# β-catenin signaling via astrocyte-encoded TCF7L2 regulates neuronal excitability and social behavior

**DOI:** 10.1101/2020.11.28.402099

**Authors:** Szewczyk Lukasz Mateusz, Lipiec Marcin Andrzej, Liszewska Ewa, Urban-Ciecko Joanna, Kondrakiewicz Ludwika, Alicja Puscian, Knapska Ewelina, Inoue Hiromi, Nowakowski Tomasz Jan, Molofsky Anna Victoria, Wiśniewska Marta Barbara

## Abstract

Astrocytes play essential roles in supporting neuronal activity and synapse formation; however, mechanisms by which these functions are regulated are unclear. The Wnt/β-catenin signaling pathway plays a crucial role in brain development and is implicated in neurodevelopmental disorders including autism spectrum disorder (ASD). We sought to investigate if some impacts of Wnt signaling are mediated via astrocytes. Here we show that the canonical Wnt/β-catenin pathway is active in postnatal cortical astrocytes and that its effector, the transcription factor TCF7L2 –is expressed in astrocyte lineage cells during embryonic and postnatal development in both mouse and human. Astrocyte-specific deletion of *Tcf7l2* in the early postnatal period led to alterations in astrocyte morphology, membrane depolarization and decreased cortical neuron excitability. Mice with the conditional knockout exhibited increased sociability and social preference in a naturalistic setting. Taken together, these data reveal a key role of astrocytic Wnt signaling in shaping postnatal neuronal development and adult social behavior.

## Introduction

Astrocytes play essential roles in the regulation of neuronal development and function by promoting excitatory synapse formation, providing metabolic support to neurons, and regulating neuronal excitability^1234^. Astrocytes are specified from radial glial progenitors beginning in mid-embryogenesis^5^, but the development and functional maturation of astrocytes continues throughout the postnatal period^6^. During that period astrocytes increase in volume and process complexity, and start expressing markers of functional maturity including gap junction connexins and potassium channels^7,8^. While multiple developmental pathways, including Wnt/β-catenin signaling, coordinate astrocyte specification^9^, the roles of β-catenin cascade in the regulation of astrocyte maturation are not well understood.

The Wnt/β-catenin pathway plays multifaceted roles in brain development that varies by cell type and developmental stage^10,11,12^. In canonical Wnt signaling, β-catenin translocates to the nucleus and acts as a cofactor for the LEF1/TCF transcription factors to activates target gene expression^13^. TCF7L2 is a Wnt effector with a particularly prominent role in brain development, including promoting proliferation of radial glia^14^, driving oligodendrocyte maturation^15^ and regulation of terminal selection of thalamic neurons^16^ *TCF7L2* is expressed in both human and murine astrocyte lineage cells^17,18^ and may play a role in astrocyte specification ^19^. However, whether it plays a role in astrocyte maturation is unknown.

*TCF7L2* is a high-confidence risk gene for the development of autism spectrum disorder^202122924^ and its expression is increased in grey matter astrocytes in the cortex of autistic patients^18^. Studies in mice heterozygous for *Tcf7l2* implicated TCF7L2 deficiency in anxiety^25,26^, suggesting a role in the regulation of behavior. However, it is not known which cell types contribute to these potential effects, or how TCF7L2 impacts autism-relevant behavioral phenotypes such as social behavior and cognition.

In this study we identify an astrocyte-specific role of TCF7L2 in the regulation of maturation and brain development. Using mouse models and conditional deletion of *Tcf7l2*, we demonstrate that astrocyte-encoded TCF7L2 is required for the morphological and molecular maturation of astrocytes. We further show that the loss of *Tcf7l2* leads to decreased neuronal excitability and hypersocial behaviors in a naturalistic setting, without significantly impacting cognition. These data reveal a role of astrocytic Wnt/β-catenin signaling in restricting sociability, with implications for how human genetic variants in this pathway may affect social behavior in the context of disease.

## Results

### The canonical Wnt pathway effector TCF7L2 is expressed in maturing astrocytes

Currently available transcriptome datasets show that Wnt/β-catenin pathway effectors are expressed in fetal and adult human astrocytes suggesting that the β-catenin pathway is active in those cells^17^. To determine whether canonical Wnt pathway is active in astrocytes, we quantified nuclear localization of β-catenin protein, a hallmark of canonical Wnt signaling. At P7, nuclear β-catenin was present in more than 80% of astrocyte nuclei labeled with eGFP in the pan astrocytic reporter line *Aldh1l1-*eGFP^27^ (Figure 1A), and in less than in 30% at P60 (Supplementary Figure S1A). We further confirmed activity of the canonical Wnt pathway in flow-sorted cells isolated from the cortex of *Aldh1l1* reporter mice at E14, E17 and P7 by analyzing *Axin2* expression - a downstream marker of Wnt signaling. *Axin2* expression increased in astrocyte lineage cells during late embryonic development and into the postnatal period (Figure 1B). Nuclear localization of β-catenin and increasing expression level of the classical Wnt target *Axin2* indicate that the intracellular path of the canonical Wnt cascade is active in maturing astrocytes.

**Figure 1:**
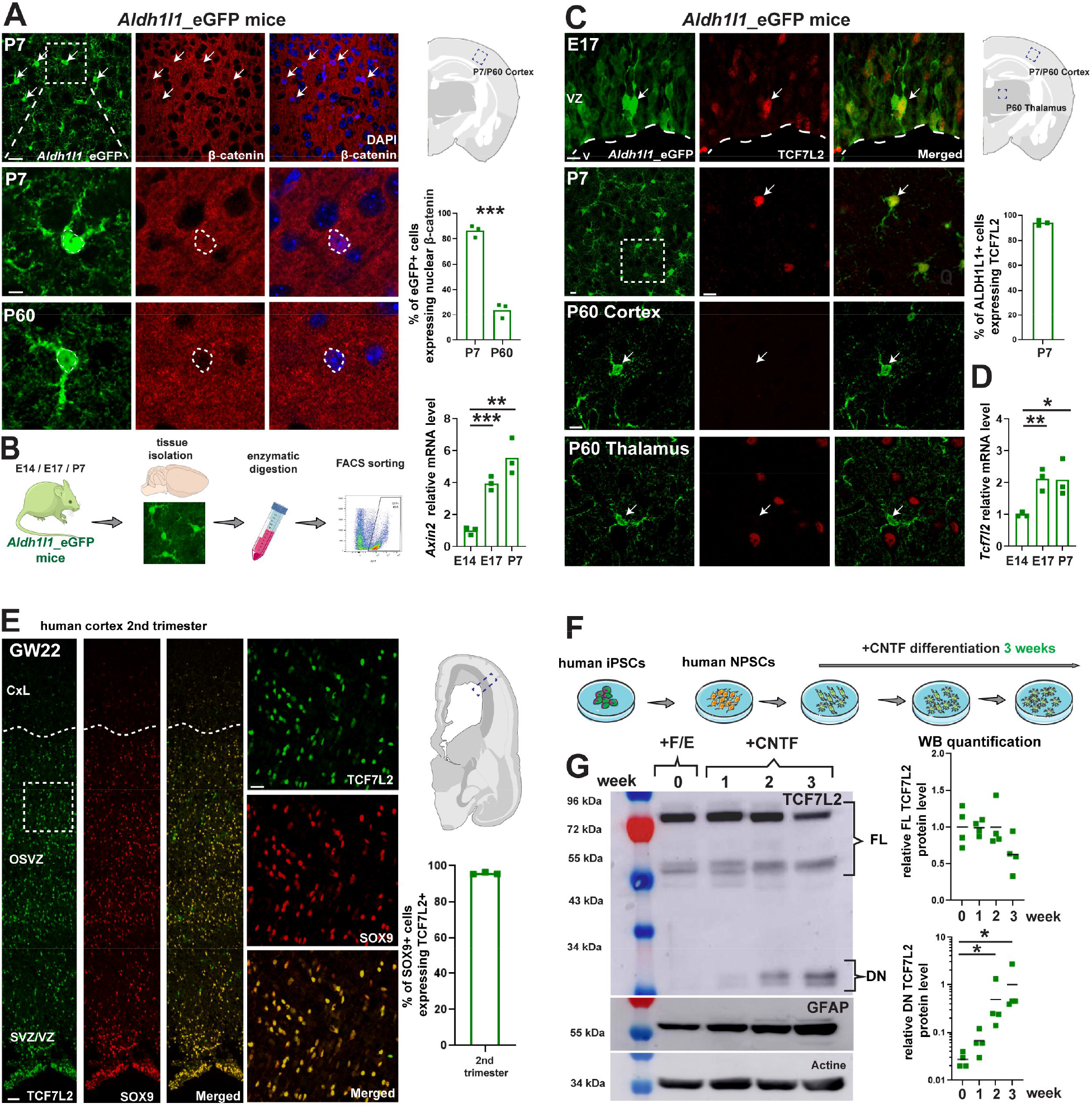
The Wnt regulator *Tcf7l2* is expressed in human and rodent astrocyte lineage cells during brain development. **A**) Images of *Aldh1l1*-eGFP+ astrocytes (green) and β-catenin (red) in the somatosensory cortex of mice on P7, n -3, scale bar – 20μm (upper), - 10μm (lower); Quantification of the number of astrocytes expressing nuclear localization of β-catenin; (**B**) Experimental workflow of isolation of eGFP-expressed astrocytes from *Aldh1l1*_eGFP mice on E14, E17, P7 using FACS sorting; RT-PCR analysis of *Axin2* expression on E14, E17 and P7 in FACS-sorted astrocytes. (**C**) Images of *Aldh1l1*-eGFP+ astrocytes expressing TCF7L2 (red) in the somatosensory cortex of mice on E17, P7 and P60, n – 3, bar – 10μm; Percentage of *Aldh1l1*-eGFP+ cells expressing TCF7L2 in the somatosensory cortex of mice on P7, n – 3; (**D**) RT-PCR analysis of *Tcf7l2* expression on E14, E17 and P7 in FACS-sorted astrocytes. (**E**) Images of TCF7L2+ cells (green) and SOX9+ cells (red) on human cortical section on gestational week 22; n – 3, scale bar - 50μm (left) - 10μm (right), Percentage of SOX9+ cells expressing TCF7L2 in the second trimester of pregnanc.; (**F**) Experimental design of *in vitro* CNTF-dependent differentiation of human NPSCs into astrocytes. (**G**) representative Western blots of TCF7L2 isoforms and GFAP in the lysates of differentiated NPSCs, w – week, densitometric analysis of TCF7L2 isoforms in human NPCs, normalized to β-actin, n – 4; SVZ/VS – Subventricular Zone/Ventricular Zone, OSVZ – outer subventricular zone, CxLay – Cortical Layers

β-catenin does not directly bind to DNA, but after the translocation to the nucleus it interacts with its effector LEF1/TCF proteins to regulate gene expression. To determine which LEF1/TCFs are expressed in astrocytes, we performed immunohistochemistry (Supplementary Figure S1B). Among all four β-catenin effectors only TCF7L2 was present in the astrocyte lineage at postnatal day 7 (Figure 1C). TCF7L2 was observed in more than 90% of astrocytes (Figure 1C, right), and about 80% of TCF7L2-positive cells were astrocytes on postnatal day 7 (Supplementary Figure 1C). Astrocytic specific expression was further confirmed in flow-sorted *Aldh1l1-*eGFP-positive cells (Figure 1D). *Tcf7l2* mRNA levels increased at late gestation, similar to *Axin2*, confirming that TCF7L2 is the major mediator of Wnt/β-catenin signaling in astrocytes. At P60, TCF7L2 was not longer observed in cortical astrocytes but was still detectable in thalamic neurons (Figure 1C, lower) and in a subset of oligodendrocyte lineage cells.

To determine if TCF7L2 is expressed also in human developing brain we immunostained slices of second trimester embryonic cortex (17-22 gestational weeks). We detected TCF7L2-positive cells throughout the ventricular and sub ventricular zones (VZ, SVZ) of which ∼ 95% co-labeled with the astrocyte marker SOX9 (Figure 1J, Supplementary Figure S2A)^17,28^. Analysis of publicly available single cell RNAseq datasets further indicated that TCF7L2 is robustly expressed in the glial lineage in the developing human brain, and is enriched in astrocytes in the adult brain (Supplementary Figure X). We also identified TCF7L2+/SOX9+ cells in VZ and SVZ of 60-day old cerebral organoids generated from human inducible pluripotent stem cells (iPSc; Supplementary Figure S2B).

Additionally, to analyze how TCF7L2 levels change during astrocyte differentiation we analyzed the expression of TCF7L2 during 3-week differentiation of human neural progenitors into astrocytes *in vitro* (Figure 1F). As astrocytes matured they expressed higher levels of GFAP and lower levels of the full-length isoform of TCF7L2 (Figure 1G). Similarly, in murine monolayer cultures, we observed that CNTF-mediated differentiation of astrocytes led to a temporal increase in *Tcf7l2* followed by a decrease in more mature cells (Supplementary Figure 2C-G). More mature human astrocytes also expressed the dominant negative isoform of TCF7L2, which was not expressed in murine astrocytes.

In summary, we found that the canonical Wnt pathway is active in developing astrocytes and that TCF7L2 is the primary Wnt effector expressed in the astrocyte lineage in both mice and humans. This raised the question of whether *Tcf7l2* plays a functional role in astrocyte maturation.

#### Tcf7l2 is cell autonomously required for astrocyte morphological and functional maturation

We next conditionally deleted *Tcf7l2* in astrocytes during the early postnatal period and examined its impact on astrocyte development. Astrocyte proliferation in rodent cortex continues into the early postnatal period^29,30^, after which they continue to develop morphologically and functionally. They increase in volume and fine process elaboration, eventually tiling throughout the gray matter^316^.

To conditionally delete *Tcf7l2* from all astrocytes, we crossed *Tcf7l2*^*tm1d/tmda*^ transgenic mice with conditional potential (*Tcf7l2*^*fl/fl*^) (Supplementary Figure 3A) with *Aldh1l1*^CRE/ERT2^ mice^32^. *Tcf7l2* knockout was induced in *Aldh1l1*^*CRE/ERT*^:*Tcf7l2*^*fl/fl*^ (cKO) mice by tamoxifen intraperitoneal injections at P6 and P8 (Figure 2A), and tamoxifen-treated *Aldh1l1*^*WT*^:*Tcf7l2*^*fl/fl*^ mice were used as controls (ctr). We observed >85% reduction of cortical astrocytes expressing TCF7L2 protein in *Tcf7l2* cKO mice relative to littermate controls (Figure 2B). We did not observe a reduction in TCF7L2 in OLIG2+ oligodendrocyte lineage cells (Supplementary Figure 3B) or in thalamic neurons (Supplementary Figure 3C), both cell types that express high levels of TCF7L2. Thus, this model efficiently and specifically led to a loss of TCF7L2 in astrocytes during postnatal development.

**Fig 2:**
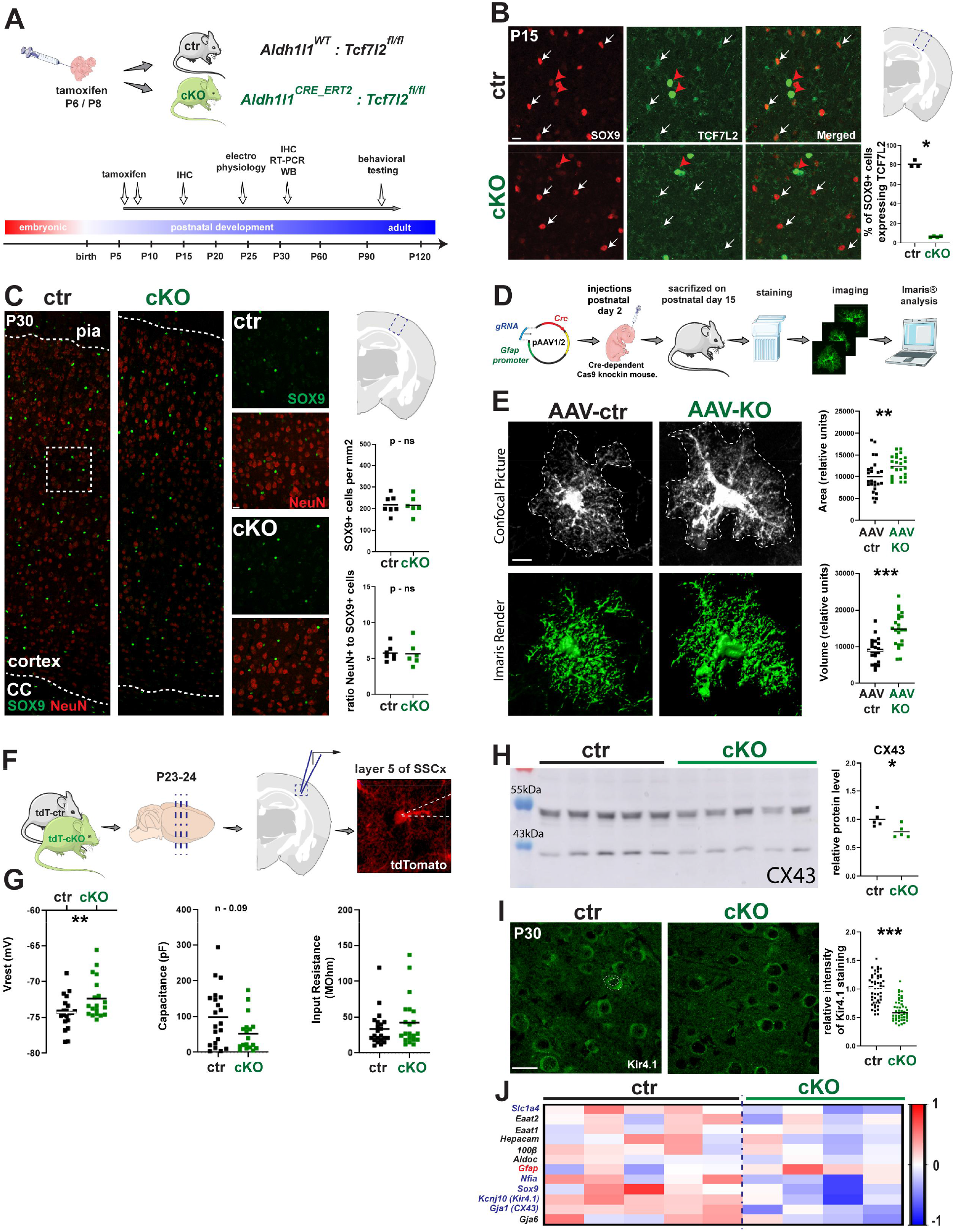
Dysmaturation of *Tcf7l2*-deficient astrocytes. (**A**) A scheme of tamoxifen-dependent *Tcf7l2* deletion in astrocytes using Aldh1l1Cre/ERT2 mice and an experimental design used in the following studies. (**B**) Images of SOX9+ (red) and TCF7L2+ (green) cells in the somatosensory cortex (SSCx) of control (ctr) and cKO mice and quantification of the effectiveness of *Tcf7l2* deletion in astrocytes at P15, n – 4, scale bar – 10μm. (**C**) Images of SOX9+ (green) and NeuN+ (red) cells in SSCx in ctr and cKO mice, n – 7, scale bar – 50μm (left),– 20μm (right) and quantification of the number of SOX9+ cells per mm2 and SOX+NeuN+ ratio. (**D**) A scheme of *in vivo* AAV-mediated *Tcf7l2* deletion in GFAP+ cells by Cas9 and workflow of analysis of morphological properties of astrocytes in vitro. (**E**) Representative images of *Gfap*-eGFP+ cells (green) in the control (AAV-ctr) and AAV-KO mice and quantification of the area and volume, n – 5, number of analyzed neurons – 22 per genotype, scale bar – 20μm. (**F**) A scheme of whole-cell patchclamp recordings from ctr and cKO tdtomato+ astrocytes. (**G**) Membrane parameters: resting potential, membrane capacitance and input resistance of ctr and cKO astrocytes; n – 19. (**H**) Representative Western blot image of Connexin 43 in the lysates of somatosensory cortex from ctr and cKO mice on P30 and their densitometric analyses, normalized to β−actin, n – 5. (**I**) images of Kir4.1 (green) cells in the somatosensory cortex of control (ctr) and cKO mice, scale bar – 10μm and semi-quantitative analysis of its protein level; n – 5, number of analyzed cells - 50 ctr and 47 cKO. (**J**) RT-PCR analysis of astrocyte markers expression in the ctr and cKO mice on P30; normalized to 18s;

We then examined the effect of *Tcf7l2* cKO on astrocyte numbers. Postnatal deletion of *Tcf7l2* in the astrocyte did not alter the density of SOX9+ cells in the cortex or the ratio of SOX9+ to NeuN+ cells (Figure 2C), suggesting that astrocyte numbers are not substantially altered by the postnatal loss of TCF7L2.

To determine cell-autonomous impacts of TCF7L2 on astrocyte morphology, we performed astrocyte sparse labeling and conditional deletion using a viral Crispr/Cas9 mediated approach. Using mice with a loxP-stop-loxP Cas9 construct (B6;129-Gt(ROSA)26Sor^tm1(CAG-cas9*,-EGFP)Fezh^/J), we injected adenovirus (AAV) expressing *Cre* under an astrocyte-specific promoter (*Gfabc1d*), as well as a guide RNA under control of a U6 promoter (Figure 2D, Supplementary Figure 4A). Efficiency of *Tcf7l2*-targeted and a control LacZ-targeted sgRNA was validated *in vitro* and in *vivo* (Supplementary Figure 4B-C). AAV was delivered by intracranial injection and astrocyte morphology was reconstructed two weeks post injection. We observed increased surface area and volume in *Tcf7l2*-deficient astrocytes relative to control astrocytes (Figure 2E). These data suggests that *Tcf7l2* restricts astrocyte process elaboration.

Astrocytes perform essential functions in supporting neuronal excitability, such as buffering of extracellular potassium and neurotransmitter reuptake^3333^. To find out whether TCF7L2 regulates basic physiologic properties of cortical astrocytes we performed whole-cell patch-clamp recordings from control and cKO layer 5 astrocytes expressing tdTomato fluorescent dye (Figure 2F). We found that *Tcf7l2* cKO astrocytes have a higher resting membrane potential compared to control cells and trended towards a lower membrane capacitance (Figure 3G). We observed decreased protein levels of the gap junctional protein connexin-43 (Figure 2H), as well as decreased expression of Kir4.1 - a potassium channel that is involved in the regulation of extracellular K^+^ concentration^33,34^ (Figure 2I). Transcripts for genes that encode these proteins, *Gja1* and *Kcna10*, respectively, were also reduced, as were several other markers associated with astrocyte functional maturation. We also observed increased expression of *Gfap*, a marker of white matter and reactive astrocytes which is not normally expressed in murine cortex (Figure 2J). Taken together, these data indicate changes in astrocyte membrane properties and molecular markers associated with a decreased capacity for gap junctional coupling and potassium reuptake.

**Fig 3.**
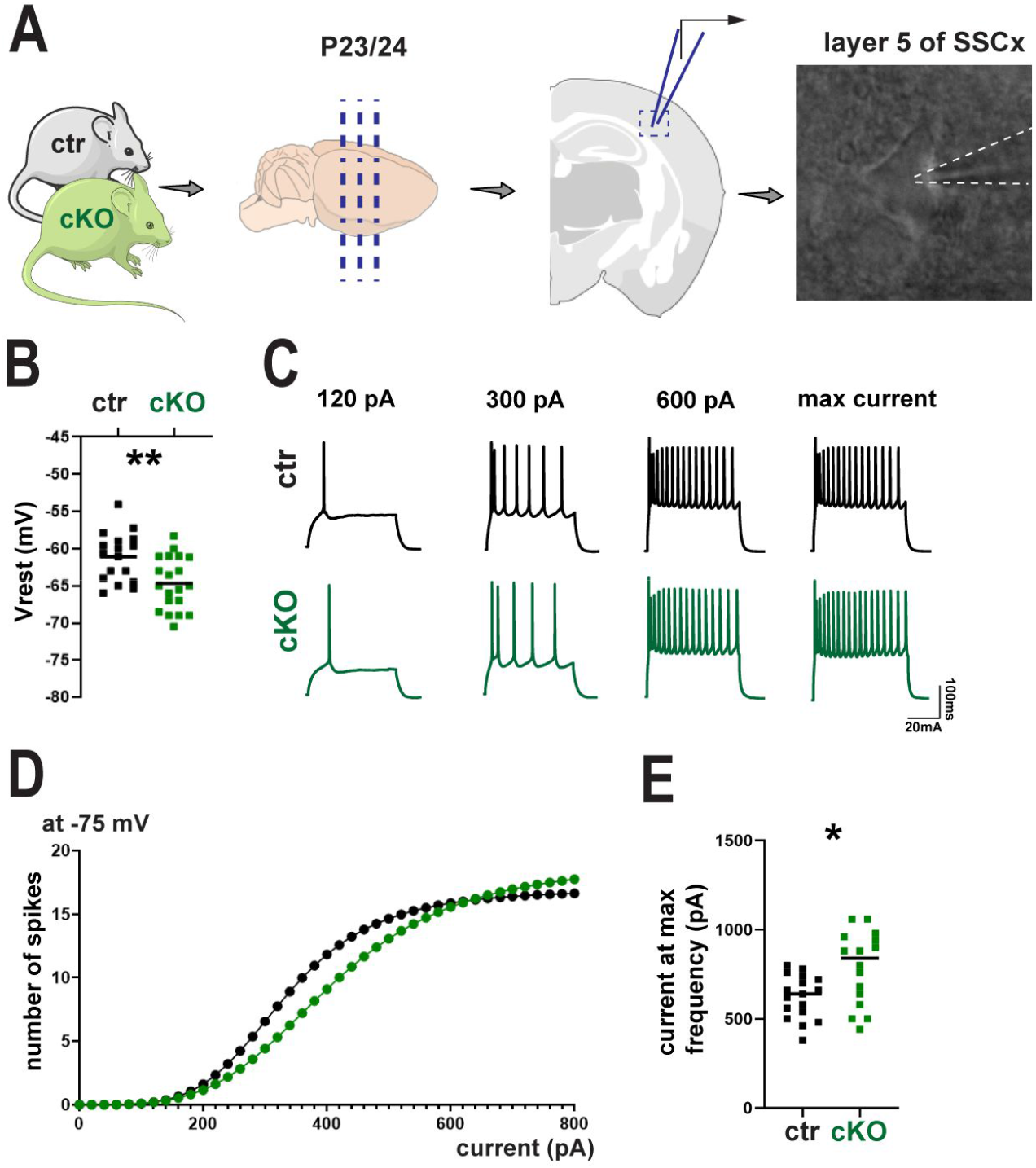
Hyperpolarized neurons in *Tcf7l2* astrocyte specific KO mice. (**A**) A scheme of whole-cell patch-clamp recordings of neurons of barrel cortex, layer 5 from control and cKO mice. (**B**) Resting potential, mV and frequency of spikes evoked by increasing depolarizing currents at −75 mV input, Hz. (**C**) Representative traces from whole-cell patch-clamp recordings of neurons in brain slices. (D) Number of spikes evoked by increasing depolarising currents at −75 mV, (**E**) Current at maximum frequency, n – 16 control, n - 17 cKO

### Astrocytic *Tcf7l2* restricts neuronal excitability

To determine whether these alterations in astrocyte morphology and function impact neuronal function, we performed whole-cell patch-clamp recordings of layer 5 pyramidal neurons in the somatosensory cortex from control and *Tcf7l2* cKO mice (Figure 4A). We found that neurons from cKO mice had a hyperpolarized resting membrane potential relative to controls, with no change in input resistance or capacitance (Figure 3B, Supplementary Figure 4D-E). Consistent with this, neurons from *Tcf7l2* cKO were less excitable, and required a higher current to reach maximal firing frequency (Figure 3C-D). Our results demonstrate that expression of *Tcf7l2* in astrocytes is required to maintain neuronal excitability, and suggest that functional deficits in astrocytes contribute to alterations in neuronal function.

**Fig 4.**
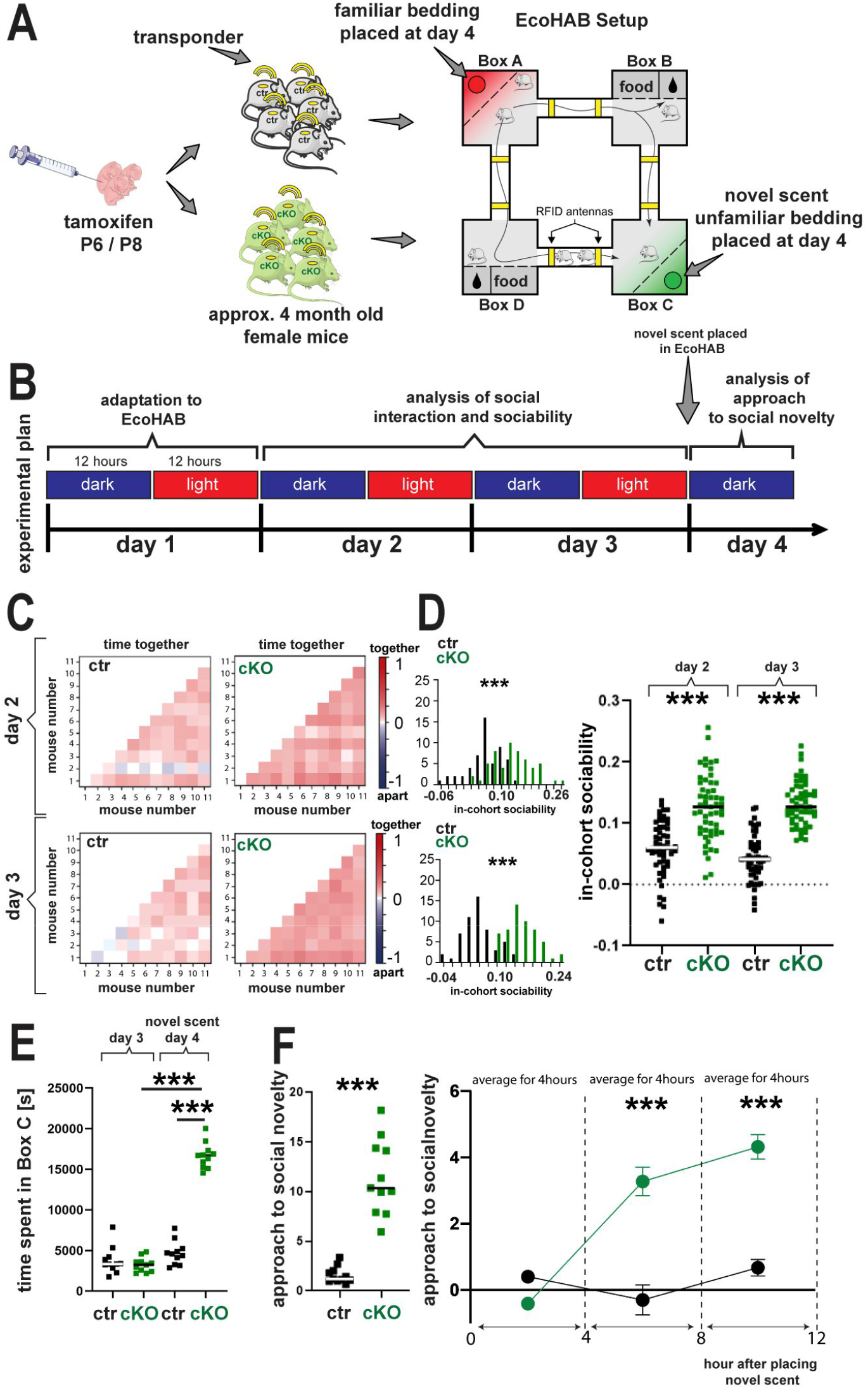
Astrocyte-specific deletion of *Tcf7l2* leads to increased sociability in a naturalistic setting. (**A**) Scheme of experimental testing of ctr and cKO mice sociability in EcoHAB system and (**B**) detailed schedule of the experiment composed of 3 phases: adaptation, sociability testing and social novelty testing. (**C**) density plot matrices for ctr and cKO cohorts on day 2 (upper) 3 (lower) of experiment;, each square represents one pair of mice and (**D**) histograms shows the distribution of this measure for all pairs of ctr and cKO mice. (**E**) Total time spent in box C (unfamiliar bedding placed on day 4) on day 3 and day 4 one dot represents one mouse. (**F**) histograms comparing approach to social novelty of ctr and cKO mice in 12h (left) or 4h bins (right), ctr n – 11, cKO n - 12

### *A*strocyte-specific depletion of *Tcf7l2* leads to hypersociability

We next asked if astrocyte-specific knockout of *Tcf7l2* impacts cognition or social behaviors. We subjected cohorts of 11-13 adult littermates of control or *Tcf7l2* cKO mice to behavioral tests in two types of naturalistic setups: EcoHAB and IntelliCage^35^. Both are computer controlled and fully automated systems used for long-term monitoring of rodent behavior, enabling objective scoring and minimizing animal-experimenter interaction^36,37^. The Eco-HAB is designed to analyze social preference and in-cohort sociability in mice^35^ while Intellicage assesses the cognitive abilities and memory^38^. All mice are tracked individually by injected transponders that emit a unique identification code when the mouse passes under RFID antennas. Intellicage and Eco-HAB thus represent a highly reproducible and naturalistic method of behavioral assessment.

We first examined social interactions using a 4-day protocol in the Eco-HAB (Figure 4A - B). After one day of habituation, we measured dynamic interactions and in-cohort sociability on days 2 and 3. *In-pair* in-cohort sociability was defined as normalized time spent with another mouse, for each pair of mice. In-cohort sociability score decreased in previously tested well-established mouse models of ASD^35^. *Tcf7l2* cKO mice displayed a two-fold increase in in-cohort sociability when compared to control mice (Figure 4C-D). Total locomotor activity was not changed (Supplementary Figure 6B). These data suggest an increased level of spontaneous social interactions.

To more specifically their responses to social novelty, on day 4 we quantified interactions with social odors after placing beddings from familiar and novel mice into the HAB. Familiar bedding (control scent) was introduced to Box A, while bedding from unfamiliar subjects (novel scent) was placed in Box C. *Tcf7l2* cKO mice spent significantly more time near the novel scent (Box C) compared to control mice (Figure 4E, left). No box preference was identified at day 3 (before placing the novel scent) suggesting that these differences were specific to the social odor (Figure 4E, right). We observed consistent results when data was plotted as the ratio of approach to the novel scent vs. control scent during the entire 12 hours after the introduction of the stimulus (Figure 4H). Taken together, our findings reveal that astrocyte-specific conditional deletion of *Tcf7l2* promotes increased social behavior.

Next we tested memory and learning in the IntelliCage setup (Supplementary Figure 5A-B). During the first three days mice were allowed to freely explore the entire setup and access all corners and chambers, which contained bottles with tap water (*simple adaptation*). After that, all corners were closed and mice learned to open a door to a corner or chamber by nosepoke (*nosepoke adaptation*). Next, the access was restricted to only one corner for each mouse (*place learning*). Visits to chambers and nosepokes were recorded for each mouse. To test learning ability, we analyzed whether and how quickly they learned to recognize the restricted corner. We calculated the number of nosepokes at the restricted corner (*correct corner nosepokes*) relative to total nosespokes. To quantify learning speed, we also analyzed the percentage of correct nosepokes after the first correct attempt. We did not identify differences between *Tcf7l2* cKO mice and littermate controls (Supplementary Figure S5C and S5D). Next, we evaluated reward motivated learning. Tap water was replaced by 10% sucrose solution (Supplementary Figure 5A). We quantified nosepokes to the sucrose-containing chamber relative to nosepokes to the tap water chamber. We observed that both control and cKO mice preferred sweetened water and that *Tcf7l2* cKO made a similar number of nosespokes to the corner with sweetened water when compared to control mice. Taken together, these data indicate that astrocytic TCF7L2 impacts social behavior without substantially impacting cognition in these assays.

## Discussion

Here we identify a novel role of the β-catenin effector TCF7L2 in regulating astrocyte maturation, and demonstrate its impact on neuronal function and social behaviour.

Our study specifically focused on the postnatal action of the β-catenin/TCF7L2 axis using temporally controlled conditional deletion to bypass impacts of Wnt/ β-catenin on early astrogenesis^19^. While both environmental and cell-autonomous processes likely contribute to astrocyte differentiation, our data show clear evidence for an instructive role of Wnt signalling through TCF7L2 in driving astrocyte maturation and function. They revealed cell-autonomous and postnatal roles for TCF7L2 in regulating astrocyte morphologic maturation and membrane properties. These data demonstrate that β-catenin signalling can play unique roles at different stages of astrocyte lineage development, as has been observed with other regulators of astrocyte development such as NFIA^39^.

We also found that astrocytes play a critical and non-neuron autonomous role in regulating neuronal function. This highlights the importance of considering cell-type specific effects when evaluating the numerous roles of Wnt signalling in synapse maturation, stability and strength^40^, including recent studies from our group showing that TCF7L2 directly regulates the expression of genes involved in neurotransmission^16^. The mechanism by which excitability of cortical neurons is decreased in mice with TCF7L2-deficient astrocytes requires further investigation. However, we did observe altered expression of astrocytic genes that are well known regulators of potassium homeostasis, including the gap-junction protein Connexin-43 and potassium channel Kir4.1^41,43^. These suggest that aberrant astrocyte maturation induced by loss of TCF2L2 could lead to altered uptake and distribution of extracellular potassium, a mechanism relevant to the regulation of neuronal excitability.

Human genetic studies predicted that *TCF7L2* is a high confidence risk gene in autism spectrum disorder^41^, that hallmark of which is impairments in social behavior. However, genomic data would seem to suggest that there are putative *loss-of-function* mutations in *TCF7L2* in ASD patients^20^. Most studies of social behavior in rodents have focused on large-effect mutations in genes like *Fmr1, Mecp2*, and others, which lead to neuronal hyperexcitability and impaired social interactions^42,43,44^.

Our study demonstrates a concordant, but inverse association, with increased sociability and decreased neuronal excitability after astrocytic knockout of *Tcf7l2*. How might this apparent discrepancy be explained? One possibility stems from our observation that human astrocytes express a TCF7L2 dominant negative isoform not present in rodent astrocytes. Depending on their impact on these different isoforms, mutations in *TCF7L2* could increase, rather than decrease Wnt signaling. Given that many ASD-associated genes can act at both ends of the sociability spectrum depending on whether they are up- or down-regulated^49^, this points to the importance of considering human-specific isoforms when interpreting behavioral data from rodent studies.

Increasing number of studies give an evidence of the involvement of astrocytes in a regulation of behaviour, such as the finding that the transcription factor MECP2 in astrocytes led to cognitive impairments in Rett syndrome^45,46,47^. Our studies establish Wnt signalling, specifically through TCF7L2 as a novel regulator of these effects. Future studies of how astrocytic dysfunction preferentially impact social behaviour could lead to new approaches to modulating astrocytes for therapeutic benefit in neurodevelopmental disorders.

## Acknowledgements

We are grateful to the Wisniewska laboratory, the Molofsky laboratory and the Knapska laboratory for the support and helpful comments on the manuscript. M.L was supported by the National Science Center - Grant OPUS 2017/25/B/NZ3/01665. The authors want to express our special gratitude to: prof. Agnieszka Kobielak, (CeNT-UW) for sharing the Ai9(RCL-tdT) mice; prof. Maciej Garstka and dr Radoslaw Mazur for help with analysis of cell morphological properties using Imaris software.

## Funding

National Science Center - Grant Sonatina 1 number 2017/24/C/NZ3/00447

## Author Contributions

LMS, AVM and MBW conceptualized the study; LMS, AVM and MBW designed experiments; LMS, MAL, EL, JUC, HI, LS performed experiments and analyzed the data; LMS created the figures; EL and TJN provided tissue; LMS acquired funding; LMS, AVM and MBW interpret the data and wrote the manuscript

## Competing Interests statement

The authors declare no competing or financial interests.

---

**Supplementary Figure 1.**
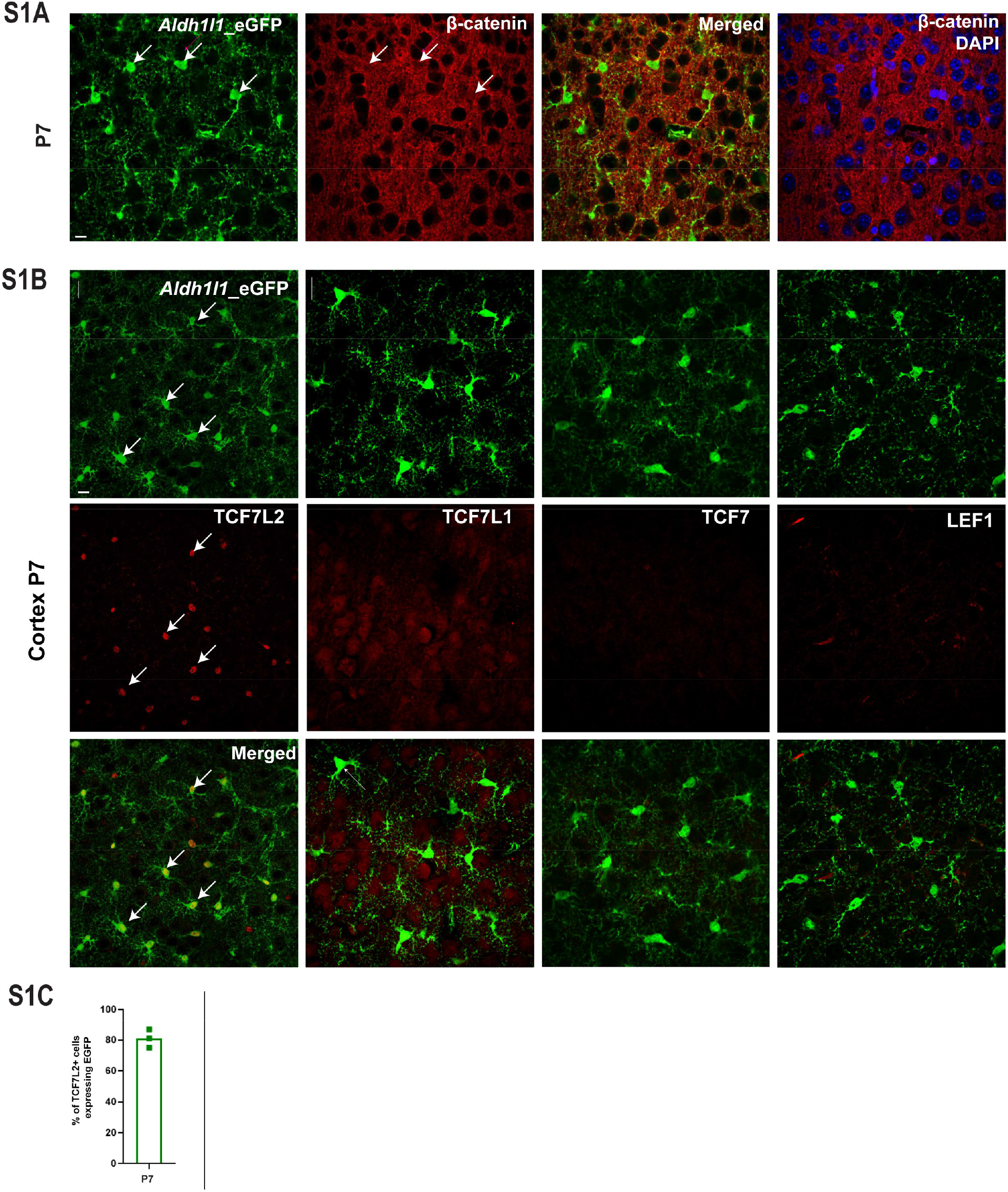
(**S1A**) Images of *Aldh1l1*-eGFP+ astrocytes (green) and β-catenin (red) in the somatosensory cortex of mice on P7 (A) and M7, n – 3, scale bar – 10μm. (**S1B**) Images of *Aldh1l1*-eGFP+ astrocytes (green) express TCF7L2 (left), TCF7L1 (left, middle) TCF7 (right, middle) and LEF1 (right) in the somatosensory cortex of mice on P7, n – 3, scale bar – 10μm (S1C) Percentage of TCF7L2+ cells expressing eGFP in the somatosensory cortex of mice on P7, n – 3

**Supplementary Figure 2.**
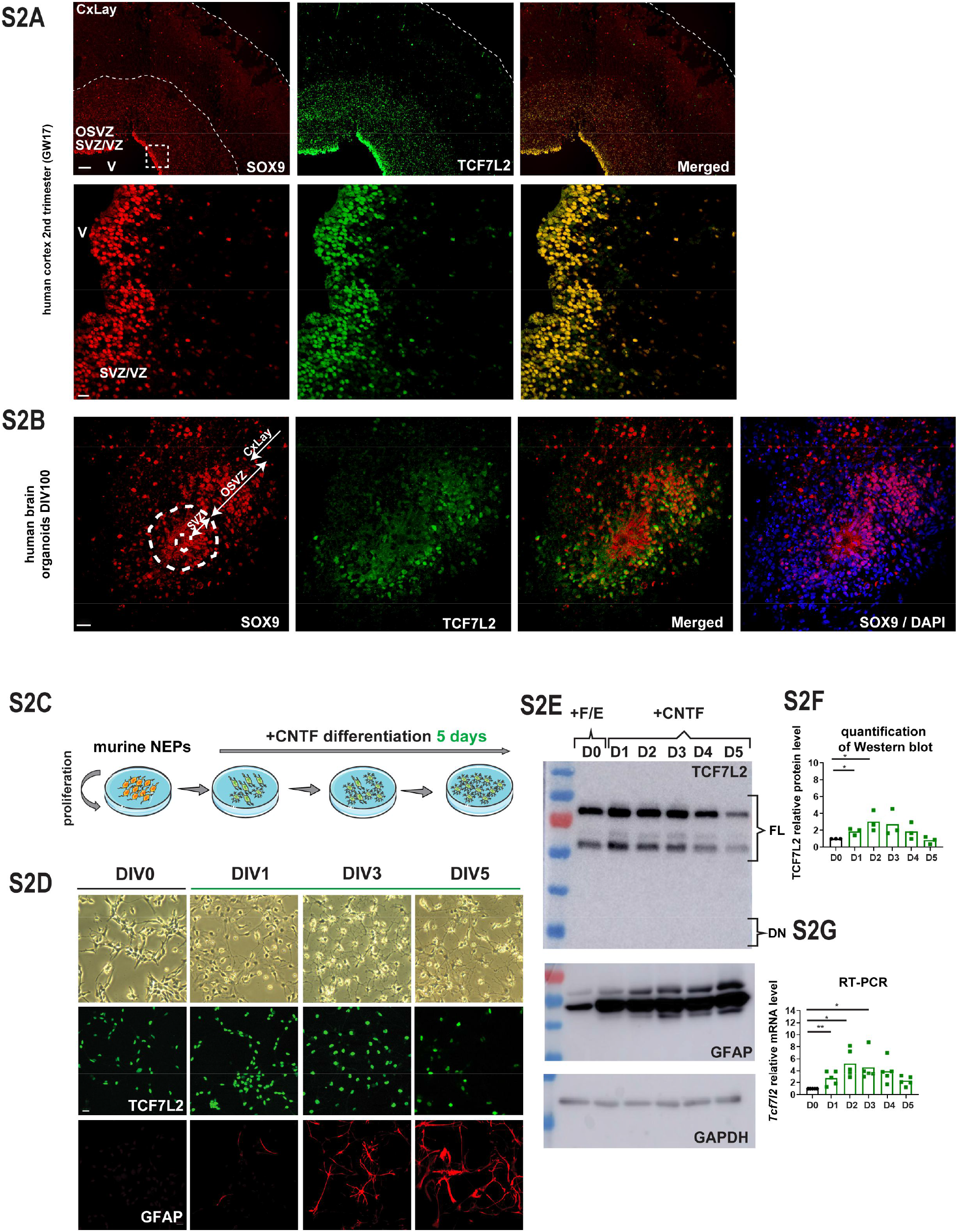
(**S2A**) Images of TCF7L2+ cells (green) and SOX9+ cells (red) on a human cortical section on gestational week 17, SVZ/VS – Subventricular Zone/Ventricular Zone, OSVZ – outer subventricular zone, CxLay – Cortical Layers, scale bar – 100μm, magnification scale bar – 20μm. (**S2B**) Images of TCF7L2+ cells (green) and SOX9+ cells (red) on 60 days old human mini-brain section SVZ/VS – Subventricular Zone/Ventricular Zone, OSVZ – outer subventricular zone, CxLay – Cortical Layers, scale bar – 20μm. (**S2C**) Experimental design for in vitro CNTF-dependent differentiation of NEPs into astrocytes. (**S2D**) Images of NEPs during differentiation; Immunostaining of TCF7L2 (green, middle row) and GFAP (red, lower row). (**S2E**) Representative western blots of TCF7L2 isoforms (upper) GFAP (middle) and GAPDH in the lysates of NEPs and densitometric analysis of TCF7L2 level, normalized to GAPDH. (**S2F**) RT-PCR analysis of Tcf7l2 expression during NEPs differentiation, normalized to Gapdh n - 5;

**Supplementary Figure 3.**
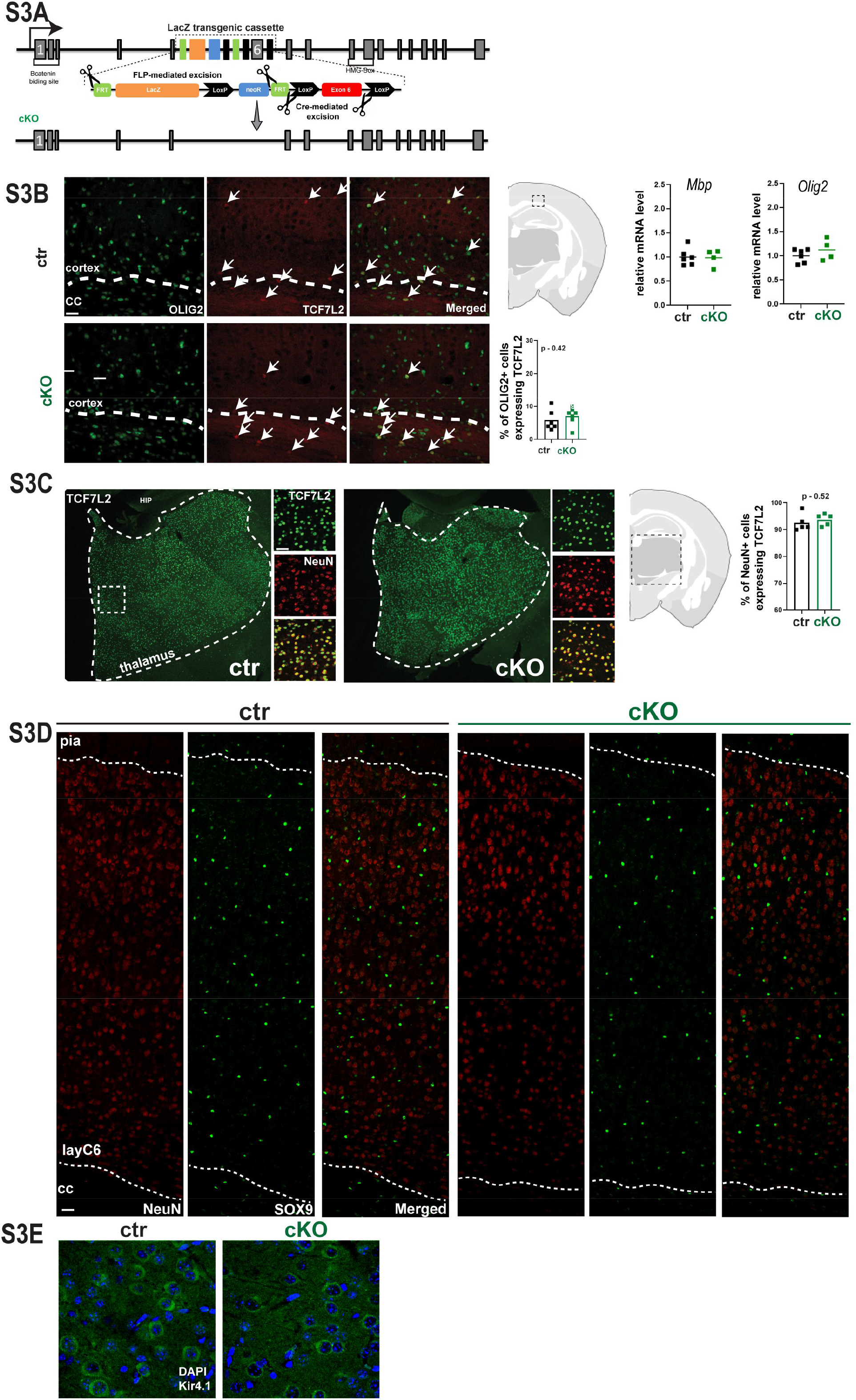
(**S3A**) Schematic representation of Tcf7l2tm1a allele generated by EUCOMM, in which a trap cassette with the lacZ and neoR elements was inserted upstream of the critical exon 6 of the Tcf7l2 gene. Exons are represented by vertical grey horizontal lines. Black arrows indicate transcription start sites. Regions that encode the β-catenin binding domain and HMG-box were also marked in the scheme of tamoxifen-dependent *Tcf7l2* deletion in astrocytes during postnatal development using of Aldh1l1Cre/ERT2 mice design. (**S3B**) Images of OLIG2+ (green) TCF7L2+ (red) cells in the somatosensory cortex of control (ctr) and cKO mice and; percentage of OLIG2+ cells expressing TCF7L2, scale bar –50μm and RT-PCR analysis of oligodendrocyte markers *Olig2* and *Mbp* in the ctr and cKO mice on P30; normalized to 18s, (**S3C**) Images of TCF7L2+ (green) NeuN+ (red) cells in the Thalamus of control (ctr) and cKO mice and percentage of NeuN+ cells expressing TCF7L2, scale bar – 100μm, magnification scale bar – 50μm. (**S3D**); Images of SOX9+ (green) and NeuN+ (red) cells in the somatosensory cortex of the ctr and cKO mice, n – 7, scale bar – 50μm, magnification– 20μm. (**S3E**) Images of Kir4.1 (green) and DAPI + cells in the somatosensory cortex of control (ctr) and cKO mice, scale bar – 10μm

**Supplementary Figure 4.**
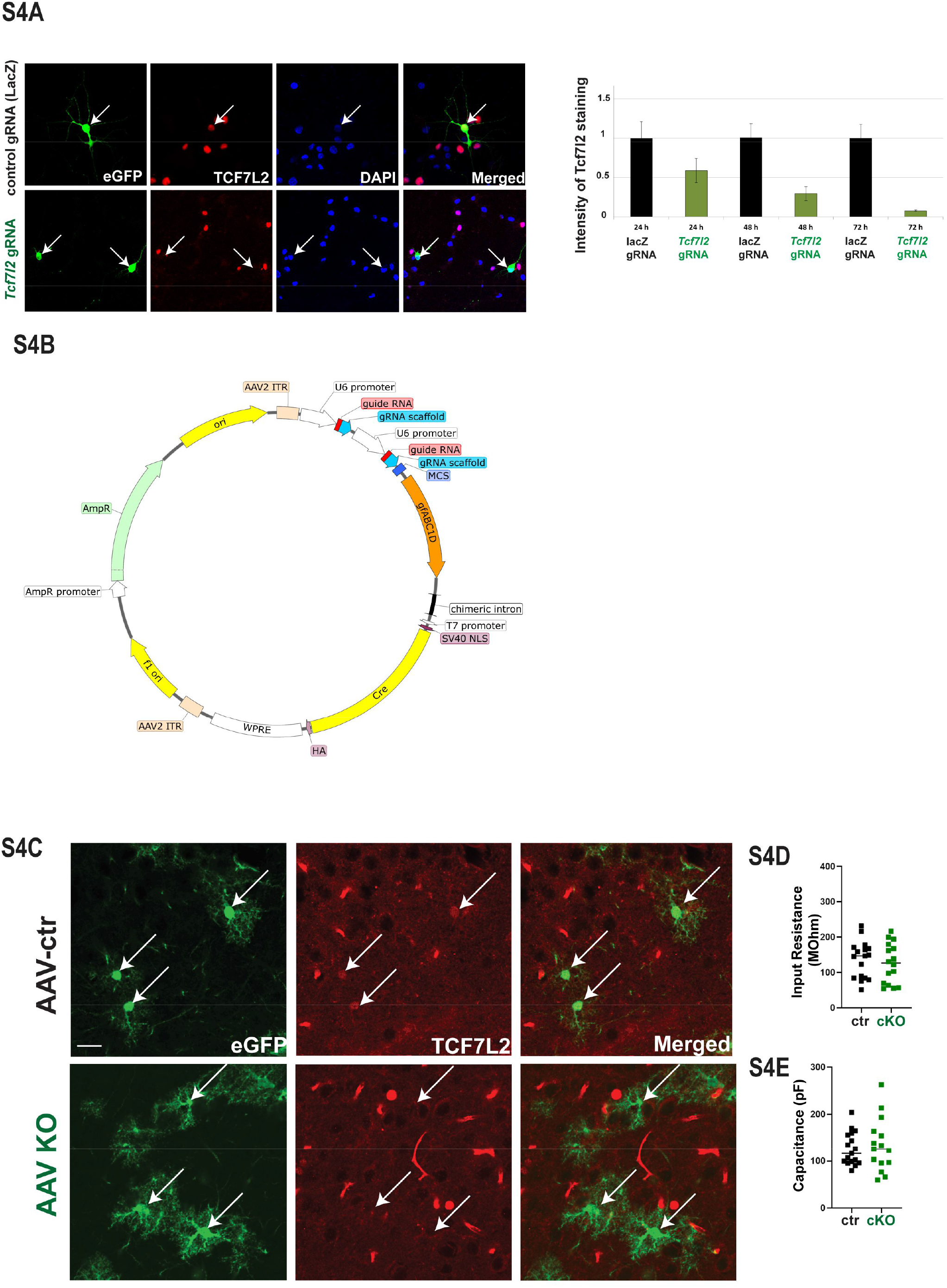
(**S4A**) Right: primary culture of thalamic neurons co-transfected with a plasmid encoding SpCas9-EGFP and a plasmid that carry gRNA against *lacZ* (control) or *Tcf7l2*. Cell culture was fixed for 24h, 48h or 72h, post-transfection and immunostained against TCF7L2 protein. Left : relative intensity of TCF7L2 fluorescence in the nuclei of eGFP-positive cells transfected with either anti-*lacZ* or anti-*Tcf7l2* gRNA. **(S4B)** Map of the gRNA-encoding plasmid used for the transfection of cell cultures and the subsequent production of AAV1/2. **(S4C)** Images of eGFP+ (green) and TCF7L2+ (red) cells in the cortex of control (AAV-ctr) and AAV-KO mice, scale bar – 20μm;

**Supplementary Figure 5.**
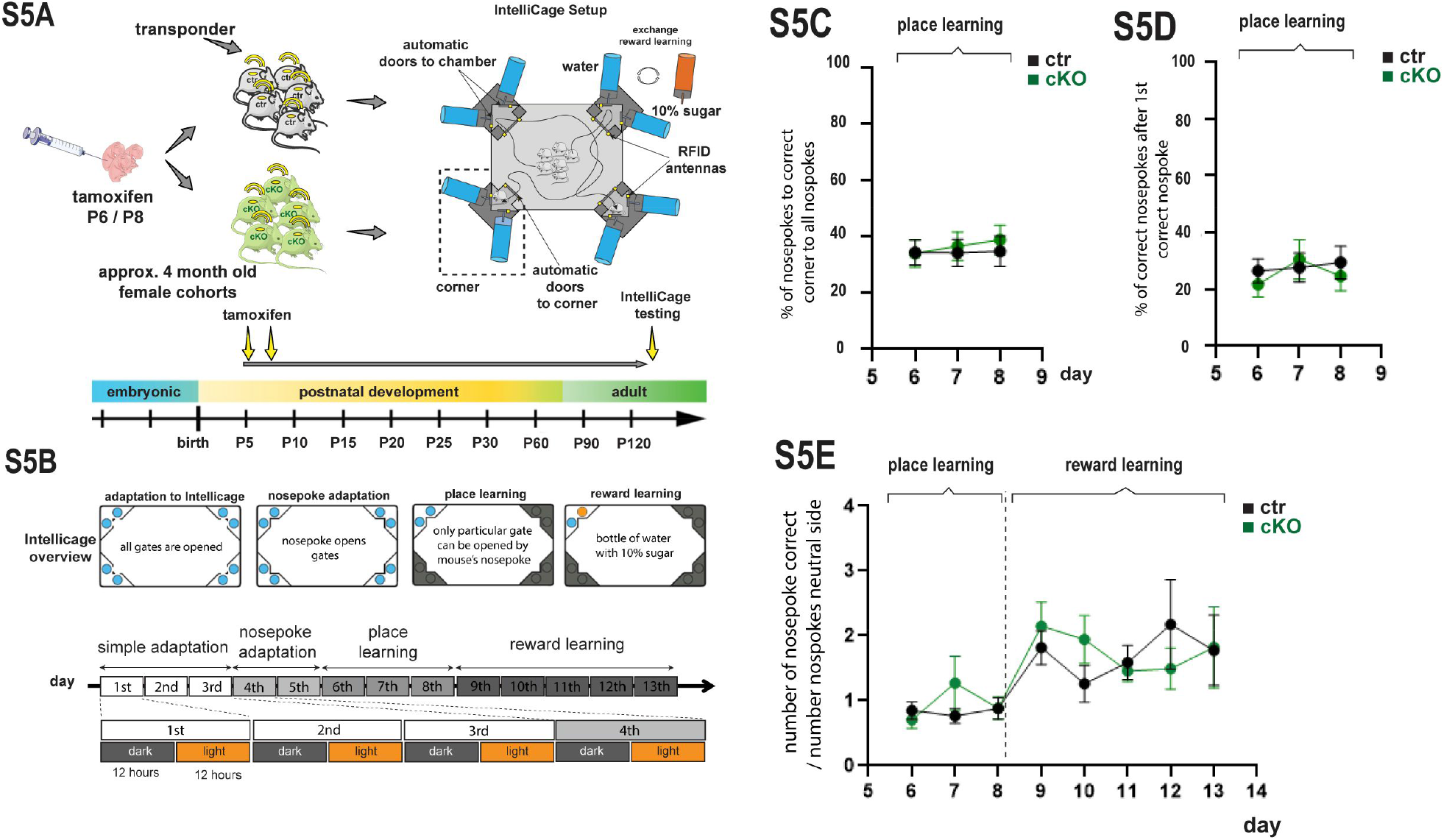
(**S5A**) A general scheme of experimental testing of ctr and cKO mice sociability in Intellicage system (**S5B**) and its experimental design and a detailed schedule of the experiment composed of 4 phases: simple adaptation, nosepoke adaptation, place preferring and place discrimination. (S5**C**) Percentage of correct nose pokes to all nose pokes. (S5**D**) Percentage of correct nose pokes after 1st correct nose poke. (S5**E**) Number of correct nose pokes to number of nose pokes to neutral side; size of cohorts: 12 control mice, 13 cKO mice

**Supplementary Figure 6.**
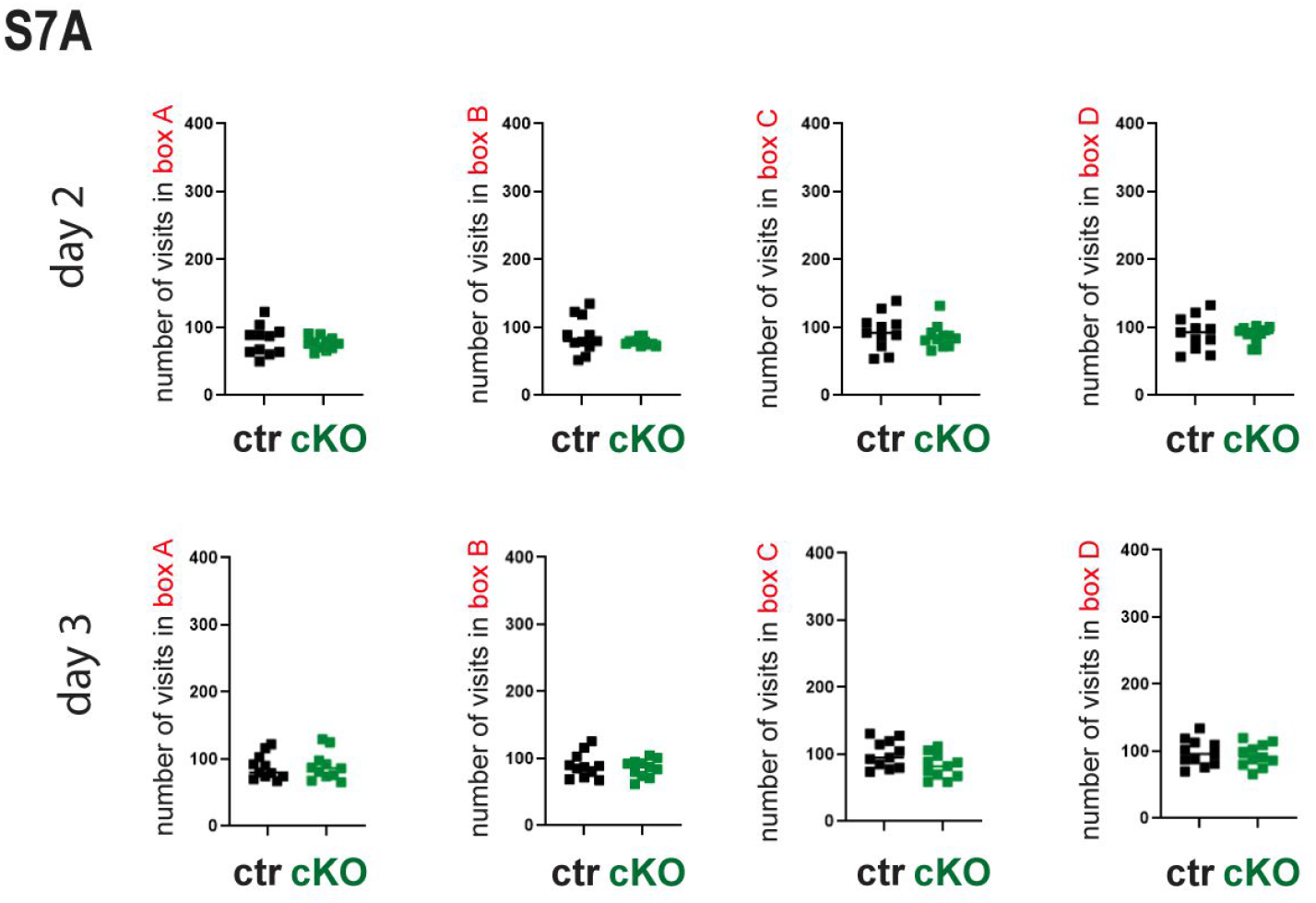
(**S6A**) Number of visits in all 4 boxes for ctr and cKO mice during 2^nd^ and 3^rd^ day of the experiment

## Materials and Methods

### Mice

We used C57BL/6NTac-Tcf7l2^tm1a^(EUCOMM)Wtsi/WtsiIeg (*Tcf7l2*^*tm1a*^) mouse strain (Skarnes et al., 2011), with a trap cassette upstream of the critical exon 6 of the *Tcf7l2* gene. Mice homozygous for the *Tcf7l2*^*tm1a*^ allele were used as total knockout of the gene. To generate the *Ald1l1*^*Cre/ERT2*^:*Tcf7l2*^*fl/fl*^ strain, in which the knockout of *Tcf7l2* is induced in the astrocytes after tamoxifen administration, *Tcf7l2*^tm1a/+^ animals were first crossed with flippase-expressing mice (ROSA26:FLPe knock-in strain; JAX stock #009086, The Jackson Laboratory^47^), and then with B6;FVB-Tg(Aldh1l1-cre/ERT2)1Khakh/J mice (Stock No: 029655, The Jackson Laboratory^32^) that express Cre recombinase from the Aldh1l1 promoter. *Aldh1l1*^*WT*^:*Tcf7l2*^*fl/fl*^ animals were used as control. To generate the *Ald1l1*^*Cre/ERT2*^:*Tcf7l2*^*fl/fl*^ :*tdTomato*^*fl*/+^ reporter strain, *Ald1l1*^*Cre/ERT2*^:*Tcf7l2*^*fl/fl*^ mice were crossed with the homozygous Ai9(RCL-tdT) strain. The *Aldh1l1*^*EGFP*^ transgenic mice, generated by the GENSAT project (The Gene Expression Nervous System Atlas (GENSAT) Project, NINDS Contracts N01NS02331 & HHSN271200723701C to The Rockefeller University (New York, NY)) encode EGFP under the control of *Aldh1l1* promoter. The B6J.129(B6N)-Gt(ROSA)26Sor^tm1(CAG-cas9*,-EGFP)Fezh^/J (*Rosa26*^*Cas9-EGFP*^) transgenic mice (The Jackson Laboratory (Stock No: 026175) (Platt et al., 2014), encode bicistronic Cas9 and EGFP cassette, whose expression under the control of CAG promoter is induced via Cre-mediated STOP-cassette removal. For the experimental procedures all mice were selected by PCR-based genotyping: *Tcf7l2tm1a* and *Tcf7l2fl* alleles: tcf_F, GGAGAGAGACGGGGTTTGTG; tcf_R, CCCACCTTTGAATGGGAGAC; floxed_PNF,ATCCGGGGGTACCGCGTCGAG;Tm1c_R, CCGCCTACTGCGACTATAGAGA; *Aldh1l1*_Cre allele: 31091, CAACAGGTGCCTTCCA; 30308,GGCAAACGGACAGAAGCA,tdTomato:oIMR9105, CTGTTCCTGTACGGCATGG; WPRE: oIMR9103, GGCATTAAAGCAGCGTATC

All animals were housed 4-5 mice/cage and maintained under standard laboratory conditions (21 ± 2 °C, humidity 60 ± 10% and 12 h/12 h dark/light cycle) with food and water provided *ad libitum*. All animals used in the present study were both males and females. All of the experimental procedures were conducted in compliance with the current normative standards of the European Community (86/609/EEC) and the Polish Government (Dz.U. 2015 poz. 266). All of the protocols for animal use were approved by the Polish Local Ethical Committee No. 1 in Warsaw. Animal usage was controlled by the institutional advisory board for animal welfare at the Centre of New Technologies.

### Neuroepithelial stem cell culture

Neuroepithelial stem cell cultures were set up according to the protocol of Hazel and Muller et al. (2001), with minor modifications. Briefly, the cortices from 3 to 4 murine E14 mouse embryos were dissected and transferred to ice-cold HBSS/HEPES. Tissue was transferred to sterile 15-ml tube and centrifuged. Pellet was resuspended in 1 ml of sterile HBSS/HEPES and dissociated, fresh 10 mL of HBSS/HEPES was added and centrifuged again. Finally, pellet containing cells was dissociated in N2 medium (DMEM/F-12 supplemented with N2 Supplement (1x), 3 g/L glucose, GlutaMAX® (1x), 100 μg/mL apotransferrin, 3mg/mL NaHCO_3_, 1% Pen/Strep) by pipetting. Cells were inoculated 1–1.5 × 10^6^ cells per dish into 10-cm dishes coated with poly-L-ornithine and fibronectin in N2 medium, and 10 ng/ml bFGF was added every second day. After 5 days of culture, cells were passaged to 6-well plates, inoculated 0.3 × 10^6^ per well in N2 medium containing 10 ng/ml bFGF. After 24h, fresh medium was added, supplemented with CNTF (10ng/uL). Cells were cultured in a medium supplemented with CNTF for 5 following days, fresh CNTF was added every day.

### Induced pluripotent stem cell reprogramming and culture

Primary fibroblasts were cultured in Dulbecco’s Modified Eagle’s Medium (DMEM) - high glucose, supplemented with 10% of fetal bovine serum and 1% penicillin-streptomycin (Sigma-Aldrich), and maintained at 37 °C in a humidified atmosphere with 5% CO_2._ At passage 4, fibroblasts were seeded at a *density* of 1 × 10^4^ *cells*/cm^2^ and on two following days transduced with lentivirus carrying Oct4, Klf4, Sox2 and c-Myc in the presence of 5ug/ml Polybrene (Sigma-Aldrich). Two days after, transduction fibroblasts were transferred onto mouse embryonic fibroblasts (MEFs) inactivated with Mitomycin C (Sigma-Aldrich) and cultured in iPSC medium composed of DMEM/F12, 20% Knockout Serum Replacement, 1% non-essential amino acid stock, 1 mM GlutaMax, 1 mM Sodium pyruvate, 100 μM β-mercaptoethanol (all ThermoFisher Scientific), 1% penicillin-streptomycin and 10 ng/mL bFGF (Alomone), with a medium change every other day. After 3 weeks, iPSC colonies were manually picked and expanded. The first 3-4 passages were performed manually every 4-5 days. Undifferentiated iPSC colonies were collected, dispersed by pipetting and small clumps of cells were plated onto fresh mitotically inactivated MEFs in iPSC medium. Subsequently, iPSC lines were adapted to feeder-free conditions and cultured on Matrigel (Corning) in Essential 8 Medium (ThermoFisher Scientific). Further, iPSCs were passaged every 4-5days using Versene solution (ThermoFisher Scientific), in the presence of ROCK inhibitor Y-27632 10uM (Tocris). The split ratio was routinely 1:6.

### Neural Induction

iPSC colonies were disaggregated using warm Accutase for 5 minutes at 37 °C and plated on Matrigel in Essential 8 Medium, in the presence of ROCK inhibitor at a density of 200 000-250 000 cells/cm^2^. The next day, the ROCK inhibitor was withdrawn and the differentiation was initiated. The cells were cultured for 4 days in KOSR medium composed of DMEM/F12, 20% Knockout Serum Replacement, 1% non-essential amino acid stock, 1 mM GlutaMax, 100 μM β-mercaptoethanol (all ThermoFisher Scientific), 1% penicillin-streptomycin supplemented with 200 nM LDN-193189, 10nM SB431542 (Tocris). At day 5, cells were passaged 1:2 and plated on Matrigel in the presence of ROCK inhibitor while the TGF-b inhibitor SB431542 was withdrawn and 200 nM LDN-193189 maintained. From day 5, increasing amounts of N2B27 media (25%, 50%, 75%, 100%) was added to the KOSR media every other day. The medium was changed daily. N2B27 medium contained DMEM/F12 : Neurobasal 50% : 50%, 1% non-essential amino acid stock, 1 mM GlutaMax, 100 μM β-mercaptoethanol, 1 x B27, 1 x N2 (all ThermoFisher Scientific), 1% penicillin-streptomycin. At day 12, the cells were passaged 1:2 and plated on Matrigel in the presence of ROCK inhibitor in NES medium: DMEM/F12, 1 mM GlutaMax, 1% penicillin-streptomycin, 1xN2, 0.05xB27, 20 ng/mL bFGF and 20 ng/mL EGF. Upon reaching confluence the neural stem cells were propagated and passaged 1:4 - 1:5 every 4-5 days using Accutase.

### Neural stem cell treatment

Neural stem cells were seeded in triplicates on Matrigel at a density of 45 000/cm^2^, in NES medium in the presence of ROCK inhibitor. The next day, the ROCK inhibitor was withdrawn and the medium was changed to the differentiation medium composed of DMEM/F12, 1 mM GlutaMax, 1% penicillin-streptomycin, 1xN2, 0.5xB27 supplemented with 10ng/ml CNTF (Alomone). The medium was changed twice a week. Samples for western blot were collected every week for 3 weeks.

### Brain organoids

The self-patterned whole-brain *organoids from iPSCs were obtained according to the protocol established by Lancaster et al. 2014. The organoids were cultured for 100 days*.

### FACS purification of astrocytes

Cortices from embryos (E14.5, E17,5) and pups (P7) were dissected and dissociated with papain 20 U/mL (Worthington) for 70 minutes at 34°C. Next, both Aldh1l1-positive and Aldh1l1 -negative cells were sorted using BD Facs Aria II and gated on forward/side scatter, live/dead by DAPI exclusion and GFP, using GFP-negative and DAPI-negative controls to set gates for each experiment. GFP-positive populations were re-sorted using the same gates. Cells were next lysed (approx. 100,000 cells per sample) using TRI-ZOL reagent (Invitrogen). Next, samples were DNAsed and purified using the RNAeasy Kit (Qiagen).

### Human Tissue Samples

De-identified fetal cortical tissue samples were collected with previous patient consent in strict observance of the legal and institutional ethical regulations from elective pregnancy termination specimens at San Francisco General Hospital. Protocols were approved by the Human Gamete, Embryo and Stem Cell Research Committee (institutional review board) at the, University of California, San Francisco

### Brain Fixing

Embryos were collected on E17.5. Timed-pregnant females were sacrificed by cervical dislocation, embryos were removed and immediately decapitated. Brains were dissected out, transferred to 4% paraformaldehyde (PFA; P6148, Sigma-Aldrich) and fixed overnight in 0.1 M phosphate-buffered saline (pH 7.4). Mice on postnatal day 8 (P8) and older were anesthetized with ketamine xylazine solution and transcardially perfused with 0.1 M phosphate-buffered saline (PBS) solution, pH 7.4, followed by 4.5% paraformaldehyde in PBS. The brains were dissected, post-fixed and cryoprotected with 30% sucrose in PBS for 24 h. Next, brains were transferred to O.C.T. (4583, Sakura Tissue-Tek) and frozen in −30°C isopentane. Sections were obtained using a Leica CM1860 cryostat. Embryonic sections were mounted directly on SuperFrost-plus slides (J1800AMNZ, Menzel-Gläser) while adult tissue was collected as free-floating sections into an anti-freeze solution (30% sucrose/30% glycerol in PBS).

### Immunocytochemistry

NEPs cells or immunopanned astrocytes cultured on coverslips were fixed with 4.5% paraformaldehyde in PBS, washed with PBS, blocked in 5% donkey serum in PBST and incubated overnight with primary antibodies (anti-TCF7L2, Cell Signalling mAb #2569, anti-SOX9, AF3075-SP, anti-βcatenin sc-7199, anti-GFAP #3670). The next day, coverslips with cells were washing and they were incubated with Alexa Fluor-conjugated secondary antibodies (Donkey anti-Rabbit IgG (H+L) Highly Cross-Adsorbed Secondary Antibody, Alexa Fluor 488, A-21206; Donkey anti-Mouse IgG (H+L) Highly Cross-Adsorbed Secondary Antibody, Alexa Fluor 594, A-21203; Donkey anti-Mouse IgG (H+L) Highly Cross-Adsorbed Secondary Antibody, Alexa Fluor 555, A-31570; Donkey anti-Goat IgG (H+L) Cross-Adsorbed Secondary Antibody, DyLight 650, SA5-10089. Finally, coverslips were assembled in Fluromount G. The images were captured with or Axio Imager Z2 LSM 700 Zeiss Confocal Microscope.

### Immunohistochemistry

Murine frozen sections on slides or free-floating slices were washed with PBST (PBS + 0.2% Triton X-100) 3 times and human sections were washed with PBST (PBS + 1% Triton X-100). Antigen retrieval was performed using sodium citrate buffer (10mM Sodium Citrate, 0.05% Tween 20, pH 6.0). Next, slices were blocked in 10% donkey serum in PBST and incubated overnight with primary antibodies (TCF7L2, #2569, anti-SOX9, AF3075-SP anti-EGFP ab13970, anti-Connexin-43 188300, anti-Kir4.1 STJ97589, anti-NeuN MAB377, anti-OLIG2 ab9610). The next day, slices were washed and incubated with Alexa Fluor-conjugated secondary antibodies (Donkey anti-Rabbit IgG (H+L) Highly Cross-Adsorbed Secondary Antibody, Alexa Fluor 488, A-21206; Donkey anti-Mouse IgG (H+L) Highly Cross-Adsorbed Secondary Antibody, Alexa Fluor 594, A-21203; Donkey anti-Mouse IgG (H+L) Highly Cross-Adsorbed Secondary Antibody, Alexa Fluor 555, A-31570; Donkey anti-Goat IgG (H+L) Cross-Adsorbed Secondary Antibody, DyLight 650, SA5-10089. Finally, slices have been assembled using Fluromount G. The images were captured with or Axio Imager Z2 LSM 700 Zeiss Confocal Microscope.

### Western Blot

Proteins from NEPs, NPCs or brain structures were extracted using ice-cold RIPA buffer (50 mM Tris, pH 7.5, 150 mM NaCl, 1% NP40, 0.5% sodium deoxycholate, 0.1% sodium dodecyl sulfate, 1 mM ethylenediaminetetraacetic acid, 1 mM NaF, Complete Protease Cocktail and Phosphatase Inhibitor Cocktail 2), centrifuged, and stored at −80°C. Proteins were separated in sodium dodecyl sulfate acrylamide gels and transferred to nitrocellulose membranes. The membranes were blocked with non-fat dry milk and incubated overnight with primary antibodies (anti-Connexin-43 188300, anti-GAPDH, 25778, TCF7L2, #2569, anti-GFAP #3670, anti-HEPACAM 18177, anti-ALDH1L1 ab87117) at 4°C. After washing, the membranes were incubated with secondary antibodies (anti-Rabbit IgG – peroxidase antibody, Sigma Aldrich, A0545; anti-Mouse IgG – peroxidase antibody, Sigma Aldrich, A9044) for 2 h at room temperature. Staining was then visualized by chemiluminescence. The images were captured using an ImageQuant LAS 4000. Densitometric analyses were performed using Quantity One 1-D software (BioRad).

### RNA isolation, cDNA synthesis and RT-PCR

RNA from NEPs, NPCs or FACS-Sorted astrocytes was isolated with the RNeasy Kit (Qiagen) and transcribed to cDNA using the Transcriptor High Fidelity cDNA Synthesis Kit (Qiagen). The levels of transcripts were measured using the SYBR Green I Master Kit (Roche) and a LightCycler 480 Instrument II (Roche). Supplementary Table 1 contains a list of used primers. Primers were designed using PrimerQuestSM. RNA from the somatosensory cortex of control and cKO mice were collected on P30. Briefly, RNA was extracted using QIAzol (79306, Qiagen) and the RNeasyMini Kit (74106, Qiagen). The quality of RNA was verified using nanodrop.

### Preparation of transfer plasmids for rAAV1/2 production

pAAV:ITR-U6-sgRNA-gfaABC1D-Cre transfer plasmid was constructed on a backbone of AAV:ITR-U6-sgRNA(backbone)-hSyn-Cre-2A-EGFP-KASH-WPRE-shortPA-ITR (Addgene #60231) (Platt et al., 2014), from which 2A-EGFP-KASH sequences were removed and hSyn promoter was replaced by gfaABC1D (similar to GFAP promoter) cloned from pZac2.1 gfaABC1D-tdTomato (Addgene #44332) (Shigetomi et al., 2013). Transfer plasmid pAAV:ITR-U6-sgRNA-hSyn-Cre had only the 2A-EGFP-KASH sequence removed, allowing for guide efficiency tests to be carried out on thalamic (TCF7L2-rich) neuronal cells. To construct control plasmids, a guide sequence against *lacZ* (TGCGAATACGCCCACGCGAT) was cloned to be expressed in tandem with gRNA scaffold under the control of hU6 promoter. For the experimental plasmids, the hU6-sgRNA cassette was duplicated and a different guide sequence against the critical 6^th^ exon of the *Tcf7l2* gene (CGTCAGCTGGTAAGTGCGG; GGTGGGGGTGTTGCACCAC) was cloned into each. Guide sequences were designed with the help of online tools CHOPCHOP v3^48^ Broad Institute GPP sgRNA Designer (https://portals.broadinstitute.org/gpp/public/analysis-tools/sgrna-design) and CRISPOR^49^

### Production, purification and titration of rAAV1/2 particles for CRISPR/Cas9-mediated gene knockout

Recombinant AAV particles of a mixed 1/2 serotype (rAAV1/2) were produced according to a modified protocol^50^. In short, HEK293T cells in exponential growth phase were simultaneously transfected with pAAV1 and pAAV2 rep/cap plasmids, pDF6 helper plasmid and pAAV:ITR-U6-sgRNA(anti-*Tcf7l2*)-gfaABC1D-Cre or pAAV:ITR-U6-sgRNA(anti-*lacZ*)-gfaABC1D-Cre (control) transfer plasmid, using PEI MAX™ MW 40,000 reagent (24765-1, Polysciences). pAAV1, pAAV2 and pDF6 plasmids were kind gifts from Lukasz Swiech (Broad Institute of MIT and Harvard, Cambridge, Massachusetts, USA). After 48-72 hours the cells were collected and lysed with sodium deoxycholate (D6750, Sigma-Aldrich), free nucleic acids were digested with Benzonase® Nuclease (70746, Millipore) and leftover cellular components were discarded after centrifugation. Cell extract was then applied in a steady rate of 1 ml/min onto HiTrap Heparin HP Column (17040601, Cytiva) equilibrated with 150 mM NaCl, 20 mM Tris, pH 8.0 buffer. The column was then washed with 20mM Tris, pH 8.0 buffer with increasing NaCl content (100mM, 200 mM, 300 mM) and the viral particles were subsequently eluted using higher salt concentrations (400 mM, 450 mM, 500 mM). With the use of Amicon® Ultra-4 Centrifugal Filter NMWL 100 KDa (UFC810024, Millipore), AAV particles were concentrated and the buffer in which they were stored was exchanged for 1x PBS. The viral batch was sterilised by passing it through Nanosep MF 0,2 µm centrifugal filter (ODPTFE02C34, Pall), aliquoted, frozen in liquid nitrogen and stored in -80°C. Titer of the AAV batch was verified by qPCR according to the modified methods from (Wang et al., 2013) and Addgene (^1^). In short, AAV aliquot was DNase-treated (AM2238, ThermoFisher Scientific), denatured in 95°C and incubated with SmaI restriction enzyme (FD0663, ThermoFisher Scientific), all according to producer’s manuals. The numbers of DNA-containing viral particles was measured using the SYBR Green I Master Kit (Roche) and a LightCycler 480 Instrument II (Roche) in a series of the treated AAV dilutions, using primers against WPRE element. Series of transfer plasmid dilutions in known concentrations were used as references, each sample was in triplicate. The viral batch was used for the experiments if its concentration was above 1·10^9^ DNA-positive rAAV1/2 particles/µl.

### *In vivo* rAAV1/2 injections

The procedure was carried out on P2 *Rosa26*^*Cas9-EGFP*^ animals of both sexes. Pups were cryoanasthetised by placing in a latex glove and immersing up to the neck in crushed ice and water for 5-8 minutes. Immobile animals were placed on a cold pack and injected with 0.5 µl of rAAV1/2 solution (either anti-*lacZ* or anti-*Tcf7l2*) into the periventricular area at the level of the septum using automated injector in a stereotaxic frame. Pups were placed on a 30°C heating pad and later returned to their mother after the recovery of mobility and if no ill effects of the procedure were observed. Animals were sacrificed on P15, their brains were isolated and prepared for immunohistochemistry.

### Electrophysiology

Brain slices (300 μm thick) from control and cKO mice (four animals per genotype) of both sexes on P21-23 were prepared using an ‘along-row’ protocol in which the anterior end of the brain was cut along a 45° plane toward the midline^51^. Slices were cut, recovered and recorded at 24°C in regular artificial cerebrospinal fluid (ACSF) composed of: 119 mM NaCl, 2.5 mM KCl, 1.3 mM MgSO4, 2.5 mM CaCl2, 1 mM NaH2PO4, 26.2 mM NaHCO3, 11 mM glucose equilibrated with 95/5% O2/CO2. The somata of layer 5a control and cKO neurons and astrocytes (CTR neurons n =13-16; ctr astrocytes, cKO neurons, cKO astrocytes in the somatosensory cortex were targeted for whole-cell patch-clamp recording with borosilicate glass electrodes (resistance 4-8 MΩ). The internal solution was composed of: 125 mM potassium gluconate, 2 mM KCl, 10 mM HEPES, 0.5 mM EGTA, 4 mM MgATP and 0.3 mM NaGTP (pH 7.25-7.35; 290 mOsm). Patch-clamp recordings were collected with a Multiclamp 700B (Molecular Devices) amplifier and Digidata 1550A digitizer and pClamp10.6 (Molecular Devices). Recordings were sampled and filtered at 10 kHz. Analysis of action potentials was performed in Clampfit 10.6. Intensity to Voltage (I-V) plots were constructed from a series of current steps in 20 pA increments from −200 to 1000 pA from a holding potential of −75 mV. A sigmoid curve (hill, 3-parameter, f = a*x^b/(c^b+x^b), Sigma Plot) was fitted to I-V of individual neurons and then parameters were compared between the 2 groups of mice. Two-tailed Mann–Whitney test or unpaired Student’s t-test were used to test for the difference in electrophysiological properties (after confirming the normal distribution of the data).

### Housing conditions and electronic tagging

Animals were group-housed under a 12 h/12 h light/dark cycle with water and food provided *ad libitum*. To minimize the effect of unstable social structure at least 4 weeks before the experiment, mice were put in cohorts used in the subsequent procedures. Under brief isoflurane anaesthesia, all mice were electronically tagged by subcutaneous injection of the glass-covered microtransponders (11.5 mm length, 2.2 mm diameter, DATAMARS for IntelliCage or 9.5 mm length, 2.2 mm diameter, RFIP Ltd for Eco-HAB) for individual identification. Microtransponders emit a unique animal identification code when activated by a magnetic field of RFID antennas or a portable chip reader. After transpondering, mice were moved from the housing facilities to the experimental rooms and adapted to the shifted light/dark cycle (the dark phase shifted from 20:00 – 8:00 to 13:00 – 01:00 or 12:00 – 24:00, depending on daylight savings time). In the housing and experimental rooms, the temperature was maintained at 23-24°C, with humidity levels between 35% and 45%.

### High throughput, automated and ecologically-relevant behavioral testing

#### Cognitive assessment – reward learning - IntelliCage

12 cKO and 12 control mice (littermates) were subjected to a 13-day IntelliCage protocol in two cohorts. First, mice were adapted to the experimental environment during a ‘simple adaptation’ phase (3 days), ‘nose poke adaptation’ phase (2 days), and ‘preparatory place learning’ phase (3 days). At this time, all drinking bottles contained tap water. During ‘simple adaptation’, doors in all conditioning units (corners) were open and access to water was unrestricted. During the following ‘nose poke adaptation’, all doors were closed by default and opened when an animal put its snout (nose poke response) into one of the two holes on the operant learning chamber’s walls. The door remained open as long as the animal kept its snout in the hole, regardless of drinking behavior. During ‘preparatory place learning’, access to drinking bottles was restricted to a single conditioning unit. A chamber with access to water was assigned randomly, with no more than 3 mice drinking from the same conditioning unit. Such design has been shown to limit social modulation of learning ^52^. This phase was followed by ‘reward learning’ (5 days). The mice then had a choice between nose-poking to the bottle containing tap water or to the bottle containing the reward (10% sucrose solution) placed behind 2 opposite doors in a given conditioning chamber. Correct response was defined as a percentage of nose pokes made to the bottle containing the reward as the first choice after entering the conditioning unit. Additionally, we measured an increase (%) in undirected nose poking after the introduction of the reward into the experimental environment, as a parameter reflecting difficulty in learning reward location. Further, for the control purposes we recorded a total number of nose pokes, visits (activity).

#### Social behavior - EcoHAB

11 cKO and 11 control mice (littermates) were subjected to 72-hour Eco-HAB protocol in two cohorts. The protocol was divided into an adaptation phase (24 h), and social behavior testing (72 h, testing approach to social odor and in-cohort sociability). During all phases of the experiment mice could freely explore all compartments. During the test of social behavior, olfactory stimuli - bedding from the cage of an unfamiliar mouse of the same strain, sex and age (novel social scent) or fresh bedding (novel non-social scent) were presented. Olfactory stimuli were placed behind the perforated partitions of the opposite testing compartments. Social approach was assessed as the ratio of approach to social vs. non-social odor during the first hour after the introduction of the stimulus. The duration of the measurement was chosen based on the activity level of the investigated mouse strain as previously described ^35^. Additionally, in-cohort sociability was assessed during the 48h-testing period. For algorithms allowing for the calculation of both sociability measures in Eco-HAB see Puscian et al, 2018

#### Statistics

Statistical analyses were performed with GraphPad Prism8 software. Two-tailed Student’s t-test was used to test for significance when comparing data. Values of P < 0.05 were considered statistically significant

## Notes

### Competing Interest Statement

The authors have declared no competing interest.

